# Individual difference in addiction-like behaviors and choice between cocaine versus food in Heterogeneous Stock rats

**DOI:** 10.1101/2021.07.21.453270

**Authors:** Sharona Sedighim, Lieselot LG Carrette, Marco Venniro, Yavin Shaham, Giordano de Guglielmo, Olivier George

## Abstract

**Rationale and objectives:** Recent studies reported that when given a mutually exclusive choice between cocaine and palatable food, most rats prefer the non-drug reward over cocaine. However, these studies used rat strains with limited genetic and behavioral diversity. Here, we used a unique outbred strain of rats (Heterogeneous Stock, HS) that mimic the genetic variability of humans.

**Methods:** We first identified individual differences in addiction-like behaviors (low and high). Next, we tested choice between cocaine and palatable food using a discrete choice procedure. We characterized the individual differences using an Addiction score that incorporates key features of addiction: escalated intake, highly motivated responding (progressive ratio), and responding despite adverse consequences (footshock punishment). We assessed food vs. cocaine choice at different drug-free days (without pre-trial cocaine self administration) during acquisition of cocaine self-administration or after escalation of cocaine self-administration. We also assessed drug vs. food choice immediately after 1-, 2-, or 6-h cocaine self-administration.

**Results:** Independent of the addiction score, without pre-trial coccaine (1 or more abstinence days) HS rats strongly preferred the palatable food over cocaine, even if the food reward was delayed or its size was reduced. However, rats with high but not low addiction score modestly increased cocaine choice immediately after 1-, 2- or 6-h cocaine self-administration.

**Conclusions:** Like other strains, HS rats strongly prefer palatable food over cocaine. Individual differences in addiction score were associated with increased drug choice in the presence but not absence (abstinence) of cocaine. The HS strain may be useful in studies on mechanisms of addiction vulnerability.

## Introduction

More than 5.5 million individuals in the United States over the age of 12 use cocaine, with 1 million diagnosed with cocaine use disorder (SAMHSA 2019). Cocaine addiction is characterized by a progressive escalation of cocaine use, increased motivation to seek the drug, and compulsive drug seeking despite negative consequences (Goldstein and Volkow 2002; Kalivas et al. 2005). There are currently no FDA-approved pharmacological treatments for cocaine use disorder. One reason for the lack of novel drug treatments is that the mechanisms of individual differences in the propensity to develop mild to severe cocaine use disorder are poorly understood (Penberthy et al. 2010; Piazza and Deroche-Gamonet 2013b). Better characterization of addiction-like behaviors in animal models may provide new insights into medication development (Venniro et al. 2020).

In the present study, we used variations of three established animal models of addiction— escalation of drug self-administration (Ahmed and Koob 1998; 1999), discrete choice between the addictive drug and non-drug palatable food reward (Lenoir et al. 2013; Lenoir et al. 2007), and the DSM-IV model (Deroche-Gamonet et al. 2004; Piazza and Deroche-Gamonet 2013a) to characterize individual differences in addiction vulnerability in Heterogenous Stock (HS) rats that mimic the genetic variability of humans (Solberg Woods and Palmer 2019).

In the escalation model, rats with extended access (6 h or more per day) but not limited access (1-2 h per day) to cocaine and other drugs increase their drug self-administration over days (Ahmed et al. 2000). In the discrete choice model, rats with a history of self-administration of cocaine or other drugs strongly prefer palatable foods over the drug when both rewards are immediately available (Ahmed 2018; Banks and Negus 2017; Caprioli et al. 2015a). In the DSM-IV rat model of addiction (Deroche-Gamonet et al. 2004; Piazza and Deroche-Gamonet 2013a), rats are classified to high, medium, or low “addiction score” based on three measures: persistent drug seeking during periods of drug unavailability (extinction responding), high motivation to self-administer the drug (assessed by progressive ratio responding), and willingness to take drug despite adverse consequences (footshock punishment) (Deroche-Gamonet et al. 2004; Piazza and Deroche-Gamonet 2013a).

However, from a translational perspective, a limitation of most previous studies using the escalation model and all previous studies using the discrete choice and DSM-IV models is that they used rat strains (Sprague-Dawley, Long-Evans, Wistar) with low genetic diversity (Solberg Woods and Palmer 2019). In the present study, we used a discrete choice procedure (Caprioli et al. 2015a; Lenoir et al. 2007) to determine palatable food versus cocaine preference in Heterogenous Stock (HS) rats with a history of extended access to cocaine self-administration (6-h/day, 0.5 mg/kg/infusion) combined with behavioral characterization of individual differences in addiction vulnerability using progressive ratio responding and footshock punishment (Deroche-Gamonet et al. 2004). Based on results from recent studies (Canchy et al. 2020; Freese et al. 2018; Vandaele et al. 2016), we also assessed individual differences in the effect of delay of reward delivery on cocaine vs. food choice and the effect of presence or absence (early abstinence) of pre-trial cocaine self administration before the choice sessions. We also assessed individual differences in food reward magnitude on cocaine vs. food choice, because this parameter plays an important role in choice in rats and monkeys (Banks and Negus 2017; Chow et al. 2020; Townsend et al. 2019).

## Materials and Methods

### Subjects

Male and female HS rats were generated by the NIH using eight inbred rat strains (ACI/N, BN/SsN, BUF/N, F344/N, M520/N, MR/N, WKY/N, and WN/N) to encompass genetic diversity (Hansen and Spuhler 1984). Twelve HS rats (Male n = 6, Female n = 6), provided by Dr. Leah Solberg Woods (Wake Forest University) were group-housed two per cage on a reverse 12 h/12 h light/dark cycle (lights off at 8:00 AM) in a temperature (20-22°C) and humidity (45-55%) controlled vivarium with free access to food and tap water. Prior to surgery, the males and females weighed 270-330 g and 180-230 g, respectively. All procedures were performed in accordance with the NIH *Guide for the Care and Use of Laboratory Animals* and were approved by the Institutional Animal Care and Use Committee of the University of California.

### Intravenous Catheterization

Rats were anesthetized with vaporized Isoflurane (1-5%). Intravenous catheters were aseptically inserted into the right jugular vein using a previously described procedure (Kallupi et al. 2020). Catheter assembly consisted of an 18-cm length Micro-Renathane tubing (0.023-inch inner diameter, 0.037-inch outer diameter; Braintree Scientific) attached to a guide cannula (Plastics One), which was bent at a right angle, embedded in dental acrylic, and anchored with mesh (1 mm thick, 2 cm circle). Tubing was inserted into the vein by puncturing it with a 22-gauge needle and was secured with suture thread. The guide cannula exited through a small incision on the back, and the base was sealed with a sealed plastic tubing and metal cover cap, which allowed for sterility and protection of the catheter base. After surgery, flunixin (2.5 mg/kg, s.c.) was administered to alleviate the pain, and fefazolin (330 mg/kg, i.m.) was administered as a form of antibiotic. Rats were allowed three days for recovery prior self-administration training, where they were monitored and their catheters were flushed daily with heparinized saline (10 U/ml of heparin sodium; American Pharmaceutical Partners) in 0.9% bacteriostatic sodium chloride (Hospira) that contained 52.4 mg/0.2 ml of the antibiotic cefazolin.

### Apparatus

Self-administration was conducted using operant chambers (29 cm × 24 cm × 19.5 cm; Med Associates) enclosed within a sound-attenuating cubicle. Chambers were constructed with a transparent plastic back and front door, two opaque metal operant panels on the left and right walls, and a stainless-steel grid floor. The right panel was equipped with two discriminative stimuli; in the far back was a house light that signaled the subsequent availability of food-paired active (retractable) lever, while in the front was a red light that signaled the subsequent availability of cocaine-paired active (retractable) lever. Presses on the food lever activated the pellet dispenser and a discrete tone cue located above the lever for 20 seconds. The pellet dispenser, located on the outside of the self-administration box, released 45 mg palatable food pellets (TestDiet; Catalogue #1811155, 12.7 percent fat, 66.7 percent carbohydrate, and 20.6 percent protein) into a food receptacle located between the two right panel levers. Presses on the cocaine-paired lever activated the infusion pump, located on top of the sound-attenuating cubicle, and a discrete yellow cue light located above the lever for 20 seconds. Cocaine was delivered via a polyethylene-50 tubing, protected by a metal spring inside the operant chamber, allowing the tubing to be connected to the rat’s IV catheter. Cocaine HCl (NIDA Drug Supply Program) was dissolved in 0.9% sodium chloride (Hospira) and was administered at a dose of 500 μg/0.1 ml/kg body weight over 4.4 seconds The left panel was equipped with a non-retractable inactive lever located in the center of the panel.

### Procedures

All procedures were conducted at the onset of the rat’s dark cycle. The food self-administration and choice procedures were a modification of previous studies with rats trained to self-administer methamphetamine, heroin, or fentanyl (Caprioli et al. 2015a; Caprioli et al. 2017; Reiner et al. 2020; Venniro et al. 2017a). A schematic representation of the experimental design is presented in Figure 1.

**Figure 1.**
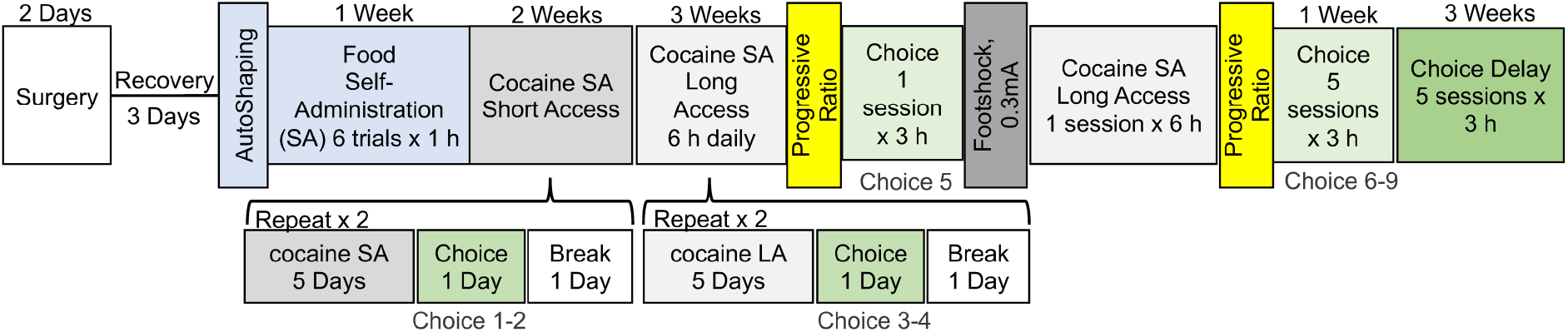
Timeline of experimental procedures. Male and female rats underwent intravenous catheterization surgery, followed by a three-day recovery period. They then underwent one food magazine session. Next, the rats self-administered the palatable food for five sessions followed by cocaine self-administration during ten 2-hour sessions (short access) and fourteen 6-hour sessions (long access). After every fifth cocaine self-administration sessions, the rats underwent a discrete choice session. Progressive ratio and footshock testing were conducted after the long-access self-administration phase, which was followed by consecutive discrete choice testing. A delay in food reward (90 s) and a decreased in the amount of food reward (from 5 pellets to 1 pellet) was used in the last five discrete choice sessions.

### Food magazine training

Rats were placed in their self-administration chambers for a single 1-hour session. Sessions began with the presentation of the house light, which remained lit for the duration of the session, serving as a discriminative stimulus for food availability. Five food pellets were released non-contingently over 5 seconds from the pellet dispenser every 10 minutes. At the end of the 1-hour session, the house light was turned off.

### Food Self-Administration

Rats were placed in their self-administration chambers for six 1-hour trials separated by a 10-minute off period, under a fixed-ratio-1 (FR1) 20-second timeout reinforcement schedule. Sessions began with the presentation of the houselight, which remained lit for the duration of the session, serving as a discriminative stimulus for food availability. Insertion of the food-paired active lever occurred 10 seconds after the houselight turned on. Food pellets were administered at a rate of 1 pellet per second for 5 seconds and food was paired with the 20-second discrete tone cue. At the end of each 1-hour session, the house light was turned off and the active lever was retracted. The rats were given 5 food self-administration sessions.

### Cocaine Self Administration

Rats were trained to self-administer cocaine during two (short access) or six (long access) 1-hour daily sessions separated by a 10-minute off period, under an FR1 20-second timeout reinforcement schedule. The daily sessions started at the onset of the dark cycle and began with the presentation of the red light, which remained lit for the duration of the session, serving as a discriminative stimulus for cocaine availability. Insertion of the cocaine-paired active lever occurred 10 seconds after the red light turned on. Pressing on the cocaine-paired lever led to the delivery of cocaine infusion (0.5 mg/kg body weight/infusion, 0.1 ml) and the 20-second discrete white light cue. At the end of each 1-hour session, the red light was turned off and the cocaine-paired active lever was retracted. During each 1-hour session, there was a limit of 15 infusions (30 infusions during short access, 90 infusions during long access). The rats were given 10 short access and 14 long access sessions.

### Progressive Ratio Testing

Progressive ratio (PR) testing was conducted with the same parameters as the cocaine self-administration sessions, including the cocaine dose and the stimulus associated with the cocaine-paired active lever. The fixed ratio number of responses per infusions were increased during this session according to the following sequence: 1, 2, 4, 6, 9, 12, 15, 20, 25, 32, 40, etc (Richardson and Roberts 1996). The breakpoint was defined as the final completed response ratio attained by the rat prior to a 60-minute period during which a fixed ratio requirement was not completed. Two progressive ratio tests were done, one following the completion of long-access cocaine self-administration and the other following punishment testing.

### Footshock Punishment Testing

A 1-hour footshock punishment testing was conducted between progressive ratio tests following the same parameters as the cocaine self-administration session. Punishment testing followed the same FR1 20-second timeout reinforcement schedule with contingent footshock (0.3 mA, 0.5 sec) paired with 30% of the cocaine infusions.

### Discrete Trials Choice Procedure

Parameters for these sessions (i.e. dose of cocaine, palatable food pellets per reward, stimuli associated with the two retractable levers) followed that of the food and cocaine self-administration sessions. The rats were given mutually exclusive choices (Ahmed et al. 2013) between the palatable food reward vs. cocaine infusions using a procedure based on those used in the studies of Caprioli and colleagues (Caprioli et al. 2015a; Caprioli et al. 2017; Reiner et al. 2020; Venniro et al. 2017a). Twenty 6-minute trials were run with a 3-minute off-period between each trial. Each trial began with the presentation of both discriminative stimuli, the houselight and the red light, followed by both the palatable food and cocaine-paired levers 10 seconds later. Rats had to select one of two levers within the 6-minute trial, during which they received the reward for the choice they made (palatable food or cocaine) with the associated cue (tone or white cue light, respectively). Reward delivery also resulted in retraction of both levers and turning off the food- and cocaine-discriminative stimuli. If a rat failed to respond within 6 minutes, no reward was given, both levers were retracted, and the discriminative stimuli were turned off.

After nine choice sessions were completed, a series of choice sessions was run with an increasing delay in the release of the food reward from 0 to 120 seconds in 7 steps (5 s, 15 s, 30 s, 45 s, 60 s, 90 s and 120 s). Each delay was assessed in two sessions. Following this series of tests, the rats were re-baselined for their cocaine self-administration with a 6-h session followed by a discrete choice session using a 90-seconds food delay and a decreased amount of food reward (from 5 pellets in 5 seconds to 1 pellet in 5 seconds), which was paired with the 20-second discrete tone cue. The effect of delay combined with different level of cocaine exposure was then tested by allowing rats to self-administer cocaine for 1 h, 2 h, or 6 h before by a discrete choice test.

### Statistical Analyses

Data were analyzed using Prism 9.0 software (GraphPad, San Diego, CA, USA). Self-administration data were analyzed using repeated-measures analysis of variance (ANOVA) followed by Bonferroni post-hoc tests when appropriate. Correlations were calculated using Pearson *r* analysis. The data are expressed as mean ± SEM unless otherwise specified. Values of p < 0.05 were considered statistically significant.

## Results

### Addiction score: Evaluation of Individual Differences in Addiction-Like Behaviors

The rats showed consistent responding for cocaine infusions during both short- and long-access self-administration sessions (Fig. 2A). During the experiment, inactive lever responding remained low, demonstrating strong lever discrimination.

**Figure 2.**
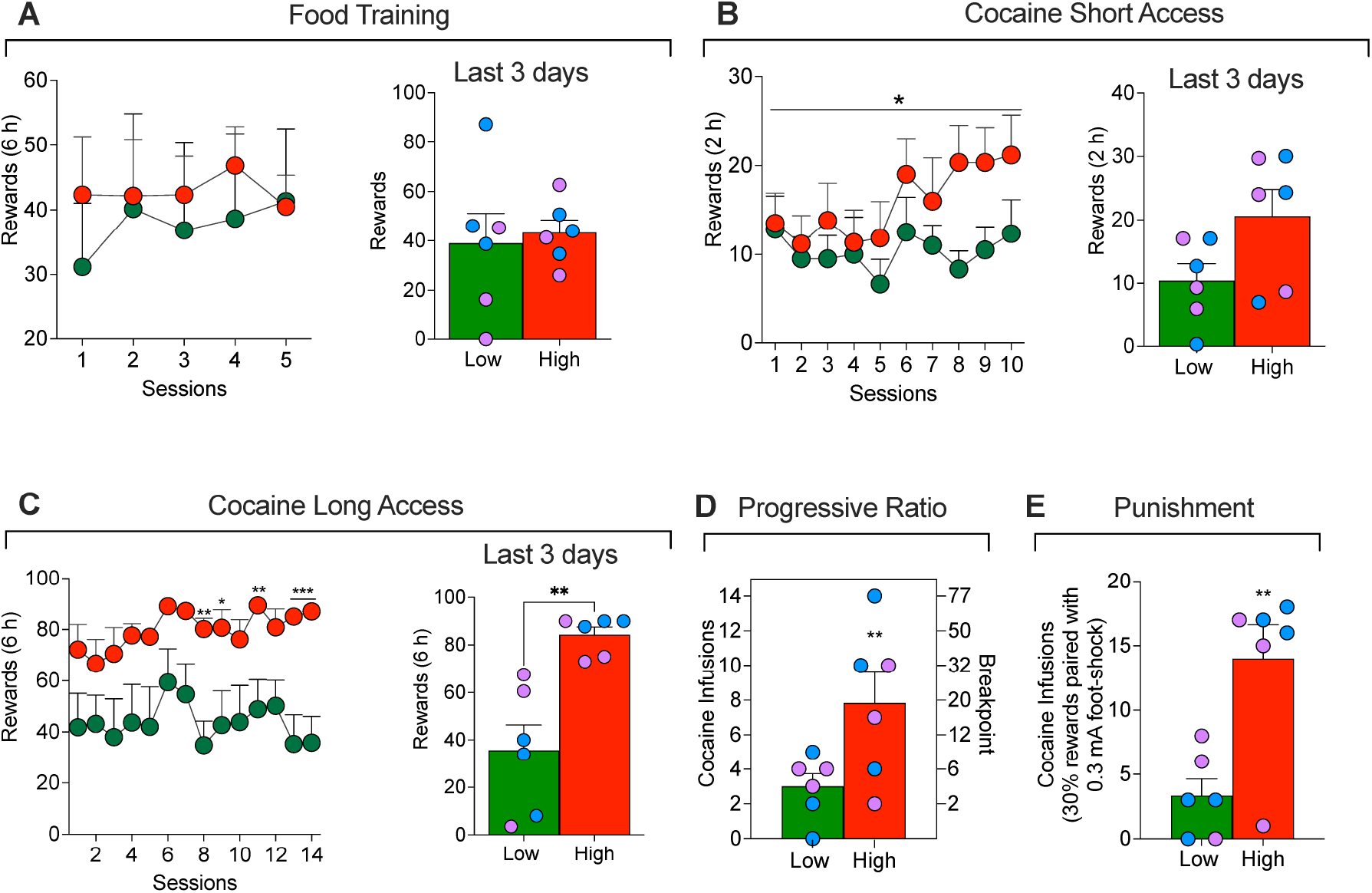
(**A**) Food training in High and Low addiction score rats. (**B**) Short-access cocaine self-administration in High and Low addiction score rats. \**p* < 0.05 *vs*. Low. (**C**) Long-access cocaine self-administration in High and Low addiction score male (blue dots) and female (red dots) rats. \**p* < 0.05, \*\**p* < 0.01, \*\*\**p* < 0.001, *vs*. A. (**D**) Cocaine self-administration on a progressive-ratio schedule of reinforcement and (**E**) and during punishment (footshock with 30% contingency) in High and Low addiction score rats. \*\**p* < 0.01, *vs*. Low.

To evaluate addiction-like behaviors, we used an Addiction score based on previous work (Deroche-Gamonet et al. 2004; Kallupi et al. 2020; Venniro et al. 2018). The score includes three addiction-like behaviors: escalation of cocaine self-administration under a fixed-ratio (FR1) reinforcement schedule, motivation to maintain responding under a progressive-ratio reinforcement schedule, and responding despite adverse consequences (footshock punishment). To calculate a composite addiction score, each measure was normalized into an index for each sex using its *Z*-score 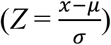, where *χ* is the raw value, *μ* is the group mean, and *σ* is the group standard deviation; the group includes both males and females (total n=12). The escalation score was calculated using the average Z-scores from the last 3 days of escalation (Fig. 2C). The progressive ratio score was calculated using the Z-score of the breakpoint value (Fig. 2D). The punishment score was calculated using the Z-scores of the cocaine rewards during footshock punishment (Fig. 2E). These scores were then combined and averaged into an addiction score.

Rats were separated into two groups, high-addicted (positive addiction score values, n=6; 3 males, 3 females) and low-addicted (negative addiction score values, n=6; 3 males, 3 females) by a median split. As expected (see Fig. 2B and 2C), significant group differences in Z-scores were seen for each individual variable: Escalation score (*t*_12_ = 4.1, *P* = 0.002), progressive ratio score (*t*_12_ = 2.3, *P* = 0.04), and punishment score (*t*_12_ = 3.6, *P* = 0.004). Additionally, the 3 addiction scores were moderately correlated: Escalation-Progressive ratio (r = 0.44, *P* = 0.14), Escalation-Punishment (r = 0.54, *P* = 0.06), and Progressive ratio-Punishment (r = 0.46, *P* = 0.13). Finally, there were no sex differences in the addiction score or any of its three measures and the data from the female and male rats were combined. Below we describe group differences between the low- and high-addicted rats during the different phases of the experiment.

### Food self-administration

The two-way ANOVA with group (low and high addicted) as the between-subjects factor and training sessions as the within-subjects factor, did not show significant differences for group (*F*_1, 10_ = 0.17, *P* = 0.6), session (*F*_4,10_ = 0.34, *P* = 0.7) and group × session (*F*_4, 40_ = 0.38, *P* = 0.8). These data demonstrate that the high and low addiction rats acquired food self-administration at the same rate (Fig 2A).

### Self-administration: short access

The two-way ANOVA with group (low and high addicted) as the between-subjects factor and training sessions as the within-subjects factor showed a significant interaction of group × session (*F*_9, 90_ = 2.1, *P* = 0.03), demonstrating that rats with high addiction score self-administered more cocaine over time, compared to low addiction score rats (Fig. 2B). However, the Bonferroni post hoc analysis did not show any statistically significant difference in any of the 10 days of short access to cocaine.

### Self-administration: long access

The two-way ANOVA with group (low and high addicted) as the between-subjects factor and training sessions as the within-subjects factor showed significant effects of group (*F*_1,10_ = 8.091, *P* < 0.05) and time (*F*_23,230_ = 41.26, *P* < 0.001) and group × time (*F*_23,230_ = 4.58, *P* < 0.001). HA rats exhibited significantly higher cocaine intake than LA rats on days 15 and 19 (*P* < 0.05), days 18 and 21 (*P* < 0.01), and days 23 and 24 (*P* < 0.001) (Fig 2C).

### Progressive ratio responding

Two progressive ratio tests were conducted during this experiment. The analysis of the combined PR data showed that high addicted rats reached significantly higher breakpoints than low addicted rats (*t*_12_ = 3.17, *P* < 0.01; Fig. 2D).

### Punishment responding

During punishment testing, the high-addicted rats showed higher resistance to punishment compared to low-addicted rats (*t*_12_ = 3.63, *P* < 0.01; Fig. 2E).

### Food vs. drug choice without pre-trial cocaine

#### Food vs. drug choice without delay

Discrete choice tests were performed after every 5 cocaine self-administration sessions (Fig. 1), as well as consecutively for 4 days after the second progressive ratio testing. A total of 9 choice sessions were completed. In order to simplify the analysis we used a *“cocaine preference score”*, defined as cocaine choices/(food choices plus cocaine choices) as the dependent measure. The Two-way ANOVA, with group (High vs Low addiction score) as the between-subjects factor and choice session as the within-subjects factor, showed significant effects of group (*F*_1,10_ = 6.2 = 0.03) and session (*F*_8,10_ = 4.7, *P* = 0.03), but no interaction (*F*_8,80_ = 1.8, *P* = 0.07) (Fig 3A). These data indicate that HS rats, independent from their addiction score or the phase of the self-administration procedure, prefer food over cocaine during the descrete choice task. This evidence is reinforced by the fact that the preference score is significantly different from the indifference level (0.5) for both groups: *t*_8_ = 22.4, *P* < 0.001 for the high addicted group and *t*_8_ = 87.8, *P* < 0.001 for the low addicted group (Fig 3A).

**Figure 3.**
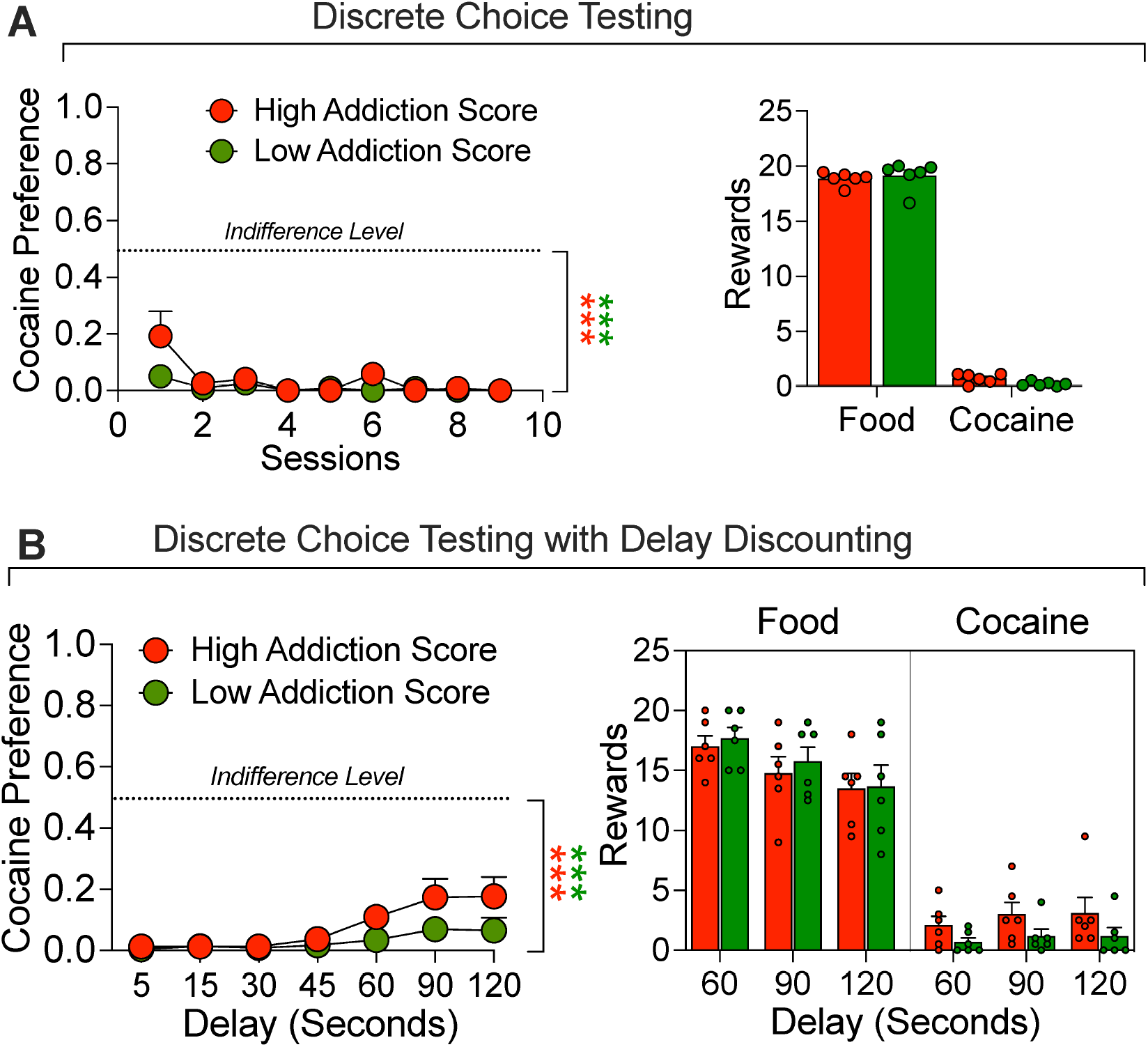
Discrete Choice Testing. **(A)** Comparison of response during discrete choice testing. All rats were given a choice between a palatable food reward or cocaine infusions for 9 test days. **(B)** Comparison of response during discrete choice testing when the food reward was delayed by 5, 15, 30, 45, 60, 90 or 120 s). The rats demonstrated high preference for food that was only modestly decreased at the long delays. \*\*\**p* < 0.001, *vs*. indifference level (0.5).

#### Food vs. drug choice with delay

After the 9 choice sessions were completed, a series of choice sessions was run with an increasing delay in the delivery of the food reward from 0 to 120 seconds in 7 steps. Each time delay time was determined in two sessions and the results were averaged. During these trials, all rats showed strong preference for food as compared to cocaine and this preference was somewhat decreased at the 90- and 120-second delays (Fig. 3*B*). The Two-way ANOVA, with group (High vs Low addiction score) as the between-subjects factor and delay as the within-subjects factor, showed a significant effect of delay (*F*_1,6_ = 7.1, *P* = 0.003) but not group (*F*_1,10_ = 2.9, *P* = 0.11) or interaction (*F*_6,60_ = 1.65, *P* = 0.11). These data indicate that HS rats, independent of their addiction score or the delay of the food reinforcer, prefer food over cocaine during the discrete choice task. This evidence is reinforced by the fact that the preference score was significantly different from the indifference level (0.5) for both groups: *t*_6_ = 14.8, *P* < 0.001 for the high addicted group and *t*_6_ = 45.9, *P* < 0.001 for the low addicted group (Fig 3B).

### Delayed (90 seconds) food vs. drug choice immediately after cocaine self-administration

#### Effect of 1-h cocaine self-administration

The effect of delay combined with cocaine exposure was then tested to further characterize differences in choice between low and high addictions score rats (Fig. 4*A*). Both groups of rats were allowed to self-administer cocaine for 1 hour under the regular training conditions, which was immediately followed by a discrete choice test. During the choice test, the food reward (5 pellets) was delayed by 90 seconds.

**Figure 4:**
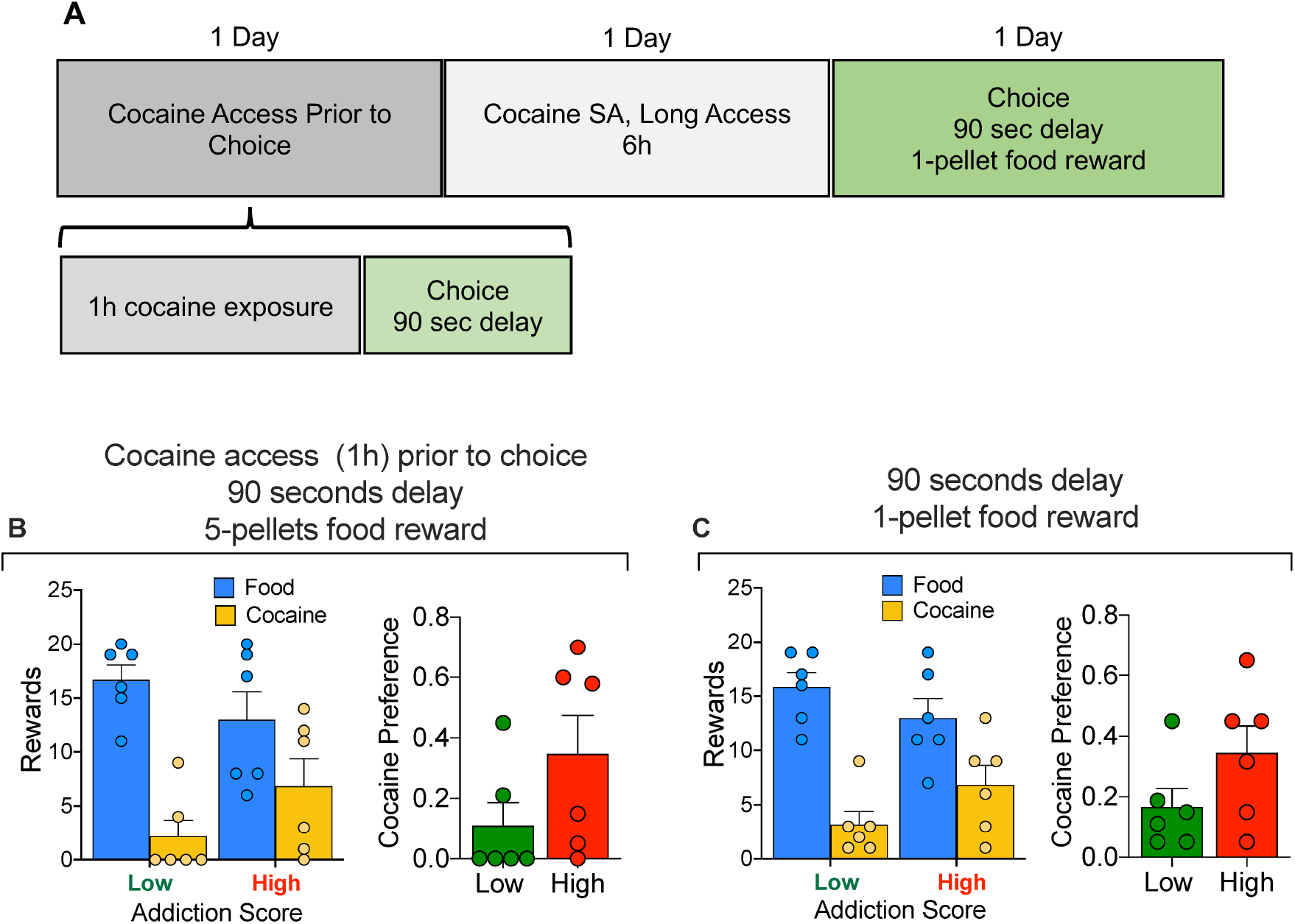
**(A)** Timeline of the experiment. **(B)** Comparison of High vs Low addiction score rats for 90-second food delay discrete choice test following 1-hour cocaine self-administration. Rats self-administered cocaine for 1 hour and were immediately tested in a discrete choice trial with a 90-second delay of the food reward (5 pellets). **(C)** Comparison of High vs Low addiction score rats for 90-second food delay discrete choice test of 1 food pellet versus cocaine. Under both conditions, the rats of both group strongly preferred the food reward

Analysis of the cocaine preference data between Low and High addiction score rats demonstrated no difference in the preference for cocaine over food between the two groups: t_10_ = 1.59, *P* = 0.14.

#### Effect of decreased food reward size

Following the choice test above and one day of 6-h cocaine self-administration, another discrete choice session was conducted; this time with both delayed (90 seconds) and decreased food reward, from 5 pellets to 1 (Fig. 4B), but without prior immediate cocaine self-administration. Food preference was preserved under these experimental conditions. Analysis of the cocaine preference data between Low and High addiction score animals demonstrated no difference in the preference for cocaine over food between the two groups: t_10_ = 1.64, *P* = 0.13.

#### Effect of 1-, 2, or 6-h cocaine self-administration

We then tested different durations of cocaine self-administration (1, 2, and 6 hours) immediately prior to a discrete choice session with the delayed (90 seconds) and reduced (1 pellet) food reward (Fig. 5A). Rats with High addiction score chose more cocaine than the low addiction score rats. The ANOVA of the cocaine preference score, which included the between-subjects factor of group and the within-subjects factor of cocaine self-administration session duration showed a significant group × session time interaction (F_2,20_ = 9.5, *P* = 0.0012). Bonferroni post-hoc tests showed higher cocaine choice in the high addiction score group in the 2-h and 6-h sessions, compared to the low addiction score group (*P* values < 0.05, Fig. 5B). Additional analysis of the first 30 minutes of the choice sessions with or without pre-trial cocaine self administration showed that the high addiction score rats, when exposed to a 6 hours pre-trial cocaine self administration session, showed increased preference for cocaine. The ANOVA of the cocaine preference score, which included the between-subjects factor of group and the within-subjects factor of cocaine self-administration session duration showed a significant group × session time interaction (F_3,30_ = 2.86, *P* = 0.05). Bonferroni post-hoc tests showed higher cocaine choice in the high addiction score group in the 6-h session, compared to the low addiction score group (*P* < 0.01, Fig. 5C). Additional analysis of this first 30-minutes bin of the choice session confirmed that only rats with low addiction score continued to show preference for food, even after pre-trial cocaine self administration (significant difference from the indifference level (0.5), *t_3_* = 17.32, *P* < 0.001, Fig. 5C).

**Figure 5.**
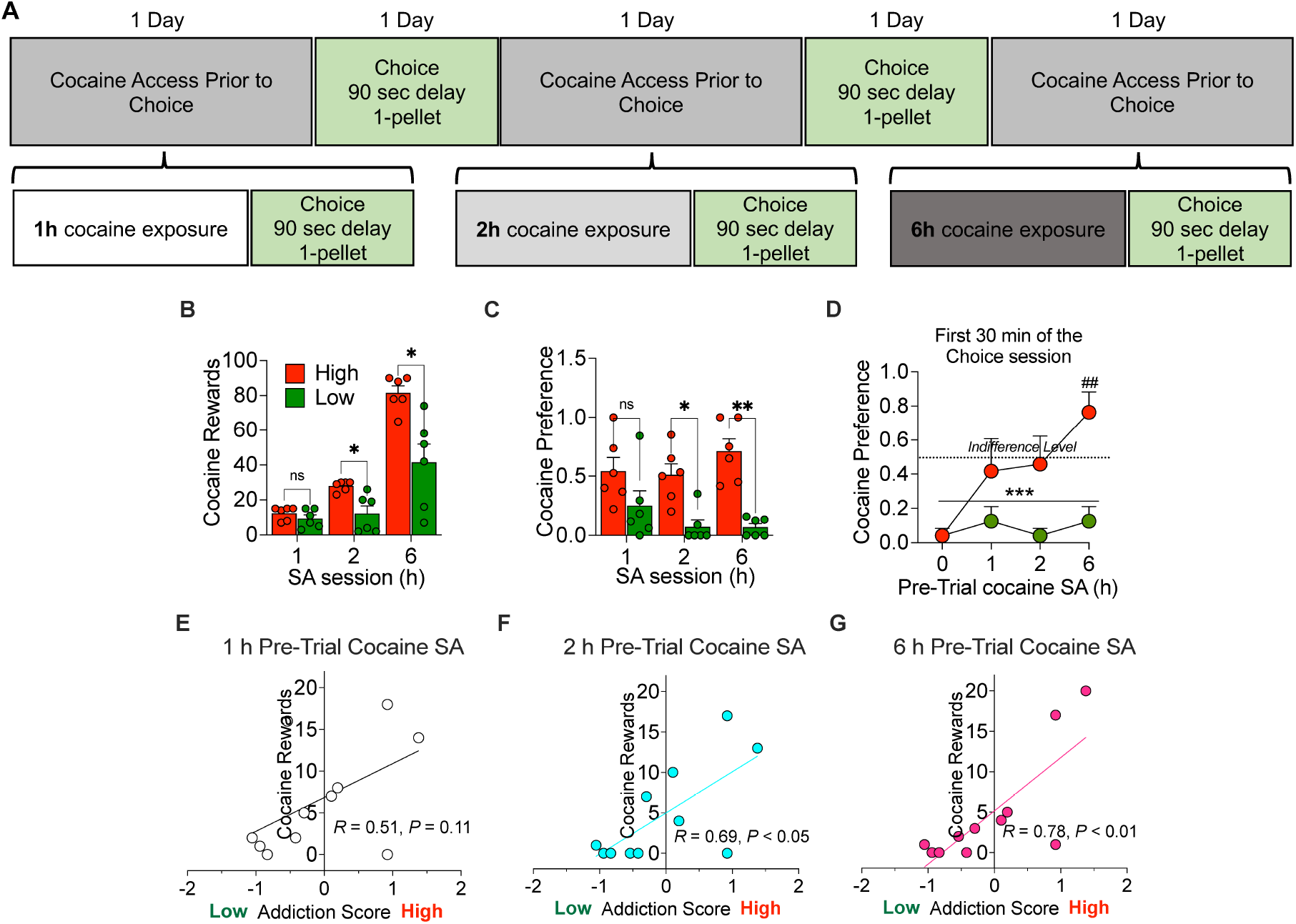
Discrete choice testing with a 90-second delay and 1 pellet food reward following 1-, 2-, and 6-h cocaine self-administration. **(A)** Timeline of the experiment. **(B)** Higher cocaine self-administration in High addicted rats than in Low addicted rats during the sessions before the discrete choice tests; * p < 0.05 vs Low rats. **(C)** High Addiction Score rats showed increased cocaine preference during the choice test that was performed immediately after a cocaine self-administration session (1, 2, or 6 hours). * p < 0.05 and ** p < 0.01 vs Low Addiction Score rats. **(D)** Analysis of the first 30 minutes of the choice test that was performed with or without pre-trial cocaine self administration. \*\*\**p* < 0.001, *vs*. indifference level (0.5), ## p < 0.01 vs Low Addiction Score rats. **(E, F, G)** Pearson correlations between the addiction score and the cocaine choice for the three duration of self-administration (*R*_1h_ = 0.51, *P* = 0.11; *R*_2h_ = 0.69, *P*_2h_ = 0.02; *R*_6h_ = 0.78, *P*_6h_ = 0.004).

Finally, correlations between the Addiction Score and the cocaine vs. food choice after different durations of cocaine self-administration are shown in Fig. 5E, F, G. The Addiction Score was significantly correlated with cocaine vs. food choice after 2-h or 6-h of cocaine self-administration (*R*_2h_ = 0.69, *P* < 0.05; *R*_6h_ = 0.78, *P* < 0.01) but not 1-h (*R*= 0.51, *P* = 0.11).

## Discussion

There are four main findings in our study. First, the HS rats showed individual differences in addiction-like behavior, as assessed in a modified version of the DSM-IV model (Deroche-Gamonet et al. 2004). Second, in absence of pre-trial cocaine self-administration, HS rats strongly preferred palatable food pellets over cocaine, even after delaying the food reward by up to 120 seconds or after decreasing food reward magnitude from 5 pellets to 1. These results extend those from previous discrete choice studies using Sprague-Dawley and Wistar rats trained to self-administer cocaine, methamphetamine, heroin, and fentanyl (Caprioli et al. 2015b; Lenoir et al. 2007; Reiner et al. 2020; Venniro et al. 2017b). Third, immediately after cocaine self-administration (intoxicated state), HS decreased their food preference. These results are in agreement with results from a previous study using Wistar rats (Vandaele et al. 2016). The fourth and most important finding in our study was that the decreased preference for delayed (90 seconds) food reward immediately after cocaine self-administration was stronger in high addiction-score rats when the food reward magnitude was decreased from 5 pellets to 1 pellets. Under these conditions, cocaine preference strongly correlated with the rats’ addiction score. Finally, in the current sample of HS rats we did not observed statistically significant sex differences in the different measures of the addiction score or the composite score. These data, however, should be interpreted with caution, because the sample size in our study was relatively small and our study was not powered to detect sex differences. Overall, our study suggests that individual difference in addiction-like behaviors in HS rats can predict increased drug preference during intoxication and that the HS rats may be useful to study behavioral and neurobiological mechanisms of individual differences in addiction vulnerability.

A critical step in the study of behavioral mechanisms of drug addiction is the use of animal models that incorporate key behavioral endpoints that are used in the diagnosis of cocaine use disorder in humans (Piazza and Deroche-Gamonet 2013a; Venniro et al. 2020). We used an animal model of extended access to cocaine self-administration combined with advanced behavioral analysis of the transition from controlled to escalated cocaine self-administration (Ahmed and Koob 1999; Deroche-Gamonet et al. 2004; George et al. 2008; Vanderschuren and Everitt 2004) in outbred HS rats. We use validated measures of escalation (self-administration under an FR reinforcement schedule), motivation to seek and take drugs (breaking point under a progressive ratio reinforcement schedule), and drug use despite adverse consequences (responding despite probabilistic (30%) footshock punishment). This behavioral approach allows for the calculation of an “addiction score” that can be used to identify rats that show high and low addiction-like behaviors. In our sample, high-addicted rats showed higher escalation of cocaine self-administration, higher breakpoints on the progressive ratio schedule, and higher resistance to punishment.

We next investigated whether the differences in cocaine addictive-like behaviors between high- and low-addicted rats are correlated with preference for cocaine vs. a non-drug reward (palatable food). We incorporate a choice procedure to our study, because drug use in humans involves choices between drug and non-alternative rewards (Ahmed et al. 2013; Banks and Negus 2017; Dunn et al. 2015; Preston et al. 2002; Silverman et al. 2012; Venniro et al. 2018).

Using a discrete choice procedure, we found that independent of the addiction score, the rats strongly preferred the palatable food over intravenous cocaine under limited (2-h/day) or extended (6-h/day) access to cocaine self-administration. The rats with the high addiction score showed a small increase in the cocaine choices compared to the low addiction score rats, but their cocaine preference was still low and significantly different from the indifference level (0.5), demonstrating a strong preference for the food reward. These effects do not seem related to an imbalance between food rewards (that had no limits) and cocaine rewards (15 infusions/h) during training. In fact, the rats averaged a total of 197 food-paired lever presses compared to 965 cocaine-paired lever presses during training, suggesting that there was no imbalance toward the food, but instead toward cocaine. These results agree with previous studies in Sprague-Dawley rats trained to self-administer methamphetamine or opioids under extended-access self-administration conditions (Caprioli et al. 2017; Caprioli et al. 2015b; Reiner et al. 2020; Venniro et al. 2017b). Introducing a delay for the food reward decreased food responses by about 30% and this effect was also largely independent of the addiction score. These results agree with previous work showing that increasing the delay for the non-drug reward decreases the preference for food or social rewards (Panlilio et al. 2017; Secci et al. 2016; Venniro et al. 2021; Venniro and Shaham 2020; Venniro et al. 2018). However, unlike the results of these previous studies, we did not observe a reversal of preference or similar (~50%) preference, even at a 120-s delay. We also found that reducing the food reward magnitude by decreasing the number of pellets from 5 to 1 had a minimal effect on food preference. These data agree with previous data showing that choice between cocaine and a non-drug alternative (saccharine or water) is insensitive to changes in the non-drug reward value (Vandaele et al. 2020; Vandaele et al. 2019). However, other studies showed that for both cocaine and fentanyl, reducing the magnitude of the food reward increases drug choice (Chow et al. 2020; Townsend et al. 2019).

A key finding of our study is that we identified individual differences in cocaine choice when the rats self-administered cocaine immediately before the food vs. cocaine choice sessions. We also found that after 6 h of cocaine self-administration cocaine choice was slightly increased in the first 30 min of the choice test in the high addiction score rats compared with 1 or 2 hours of pre-trial cocaine self-administration. This effect might be due to the fact that 6-h access produces higher self-administration levels and higher blood cocaine levels compared to limited access (700 nM vs. 500 nM, see (Ahmed et al. 2003)). Therefore, it is likely that the high addiction score rats after extended access of cocaine self administration were more intoxicated and more biased toward cocaine compared to either the low addiction rats or compared to themselves after 1-h or 2-h cocaine access. These results extend previous studies showing that non-contingent cocaine injections prior to discrete choice trials between cocaine vs. food increase cocaine preference (Freese et al. 2018; Vandaele et al. 2016). These result agree with studies in humans showing that cocaine intoxication at the moment of choice can also promote cocaine choices over other nondrug options (Donny et al. 2004; Vosburg et al. 2010). Additionally, under conditions of low food reward size (1 pellet) the increase in cocaine choice vs. food strongly correlated with the addiction score. Thus, individual differences in the addiction score were correlated with individual differences in choice in the presence (intoxication state) but not the absence of cocaine (pre-trial cocaine self administration). To the best of our knowledge, our study is the first to correlate individual differences in ‘addiction severity’ in the presence of the drug (intoxication after cocaine self-administration). In contrast, several recent studies assessed the relationship between ‘addiction severity’, assessed using measures similar to the ones used in our study, and choice of food or social interaction vs. drugs without pre-trial drug self-administration. Augier et al. (2018) reported that alcohol vs. saccharin choice correlates with progressive ratio responding and punishment sensitivity. In contrast, different measures of addiction-like behaviors did not correlate with choice between cocaine vs. food (Ahmed et al. 2013) or choice between methamphetamine vs. social interaction (Venniro et al. 2018). Taken together, the present results suggest that individual difference in addiction-like behaviors in HS rats can predict increased drug preference during intoxication.

In conclusion, we have recently shown that the characterization of individual differences in opioid addiction-like behaviors in HS rats can identify neuroadaptations that selectively contribute to these behaviors (Kallupi et al. 2020). Here, we report that HS rats with a history of extended access of cocaine self-administration prefer the palatable food reward over cocaine in a discrete choice procedure under standard conditions (no intoxication, 1-5 food pellets, delay 0-120s). However, after behavioral characterization of individual differences, high-addicted rats appear to exhibit higher cocaine choices than low addicted rats when given prior access to cocaine self-administration of at least 2 h. The results from these two studies suggest that the genetically diverse HS strain can be useful in studies on behavioral and neurobiological mechanisms of addiction vulnerability.

## Author Contributions

G.d.G and O.G. designed research; S.S. and L.L.G.C performed research; S.S., L.C., and G.d.G. analyzed data; and S.S. and G.d.G. wrote the paper. MV and YS contributed to data analysis and interpretation, and to the writeup of the paper. All of the authors critically reviewed the content and approved the final version for publication.

